## supplemental figures for "Experimental evolution for the recovery of growth loss due to genome reduction"

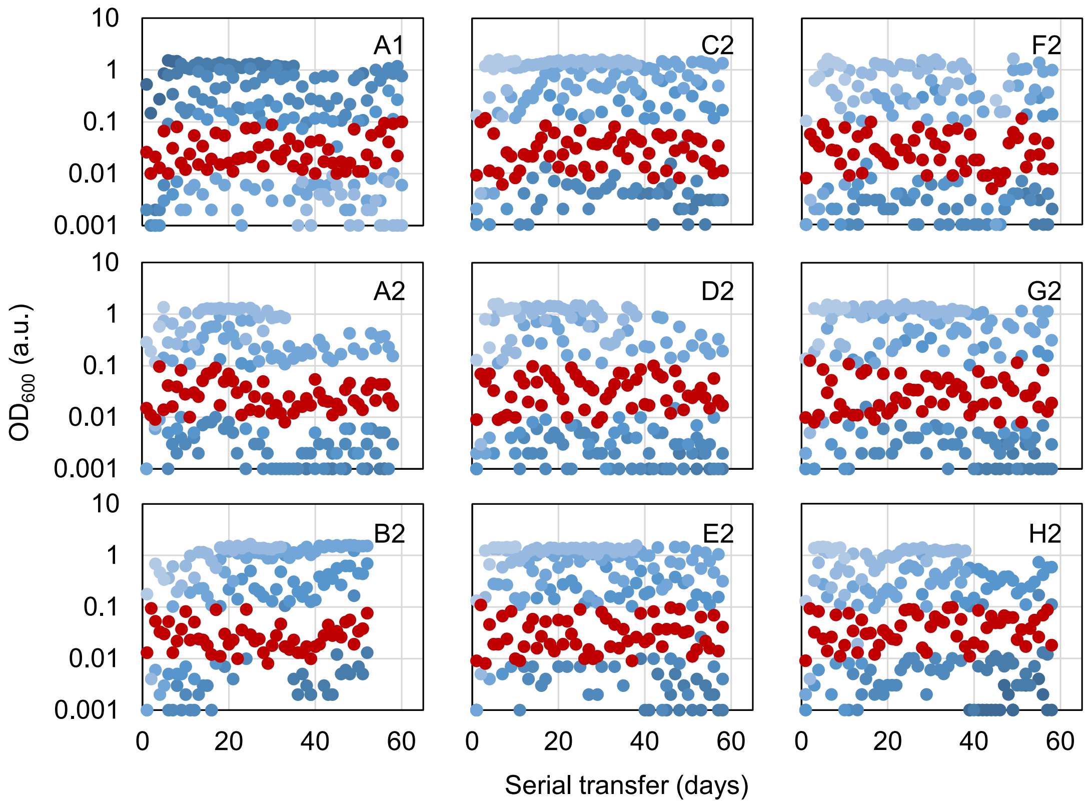


**Figure S1 Temporal $OD_{600}$ records of the overnight cultures.** Four different dilution rates of serial transfer were made for each evolutionary lineage, indicated in the blue gradation. Only one of the four overnight cultures in the exponential growth phase (OD600 = 0.01~0.1) was chosen for the following serial transfer, highlighted in red.

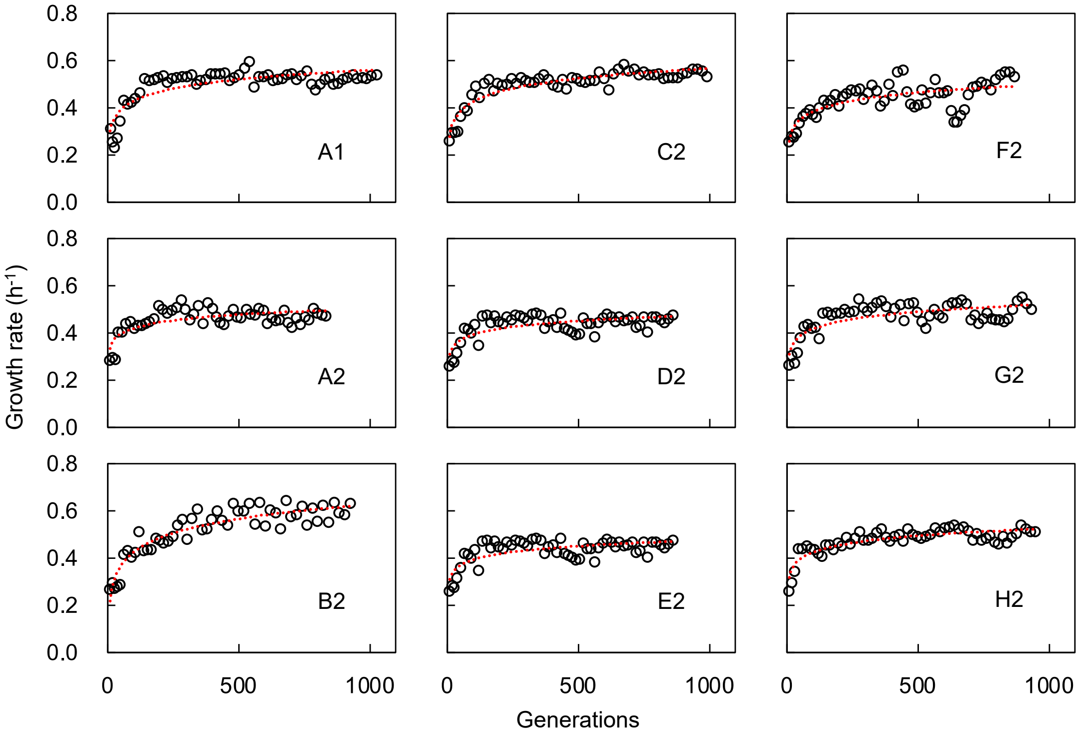


**Figure S2 Temporal changes in growth rate during experimental evolution.** The nine evolutionary lineages are shown separately, equivalent to Figure 1A. The red lines are the logarithmic regression of the changing growth rates.

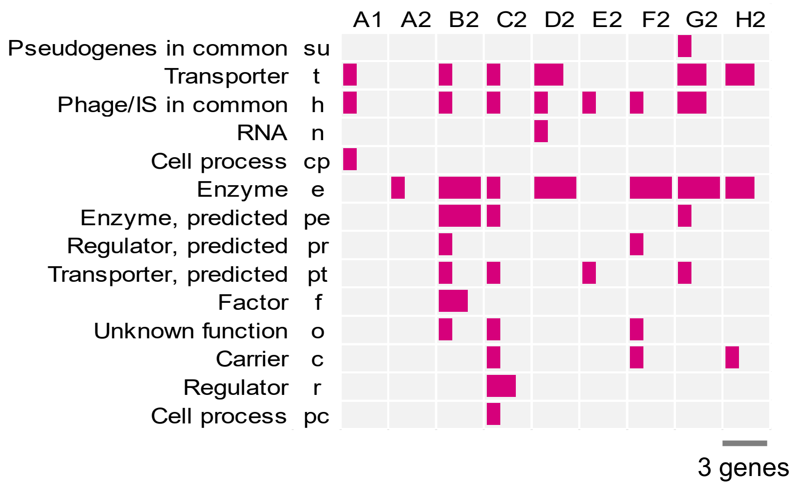


**Figure S3 Gene categories comprising the mutated genes.** The number of mutated genes assigned in the gene categories is indicated as the length of the color bar.

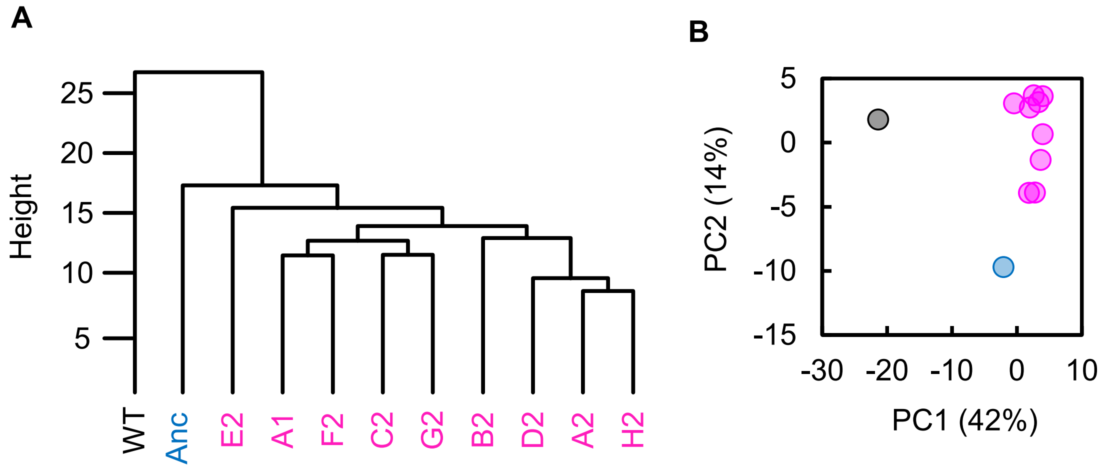


**Figure S4 Clustering and principal component analysis of transcriptomes. A.** Hierarchical clustering. **B.** Principal component analysis. PC1 and PC2 components are shown. Black, blue, and pink represent the wild-type, reduced, and evolved genomes, respectively.

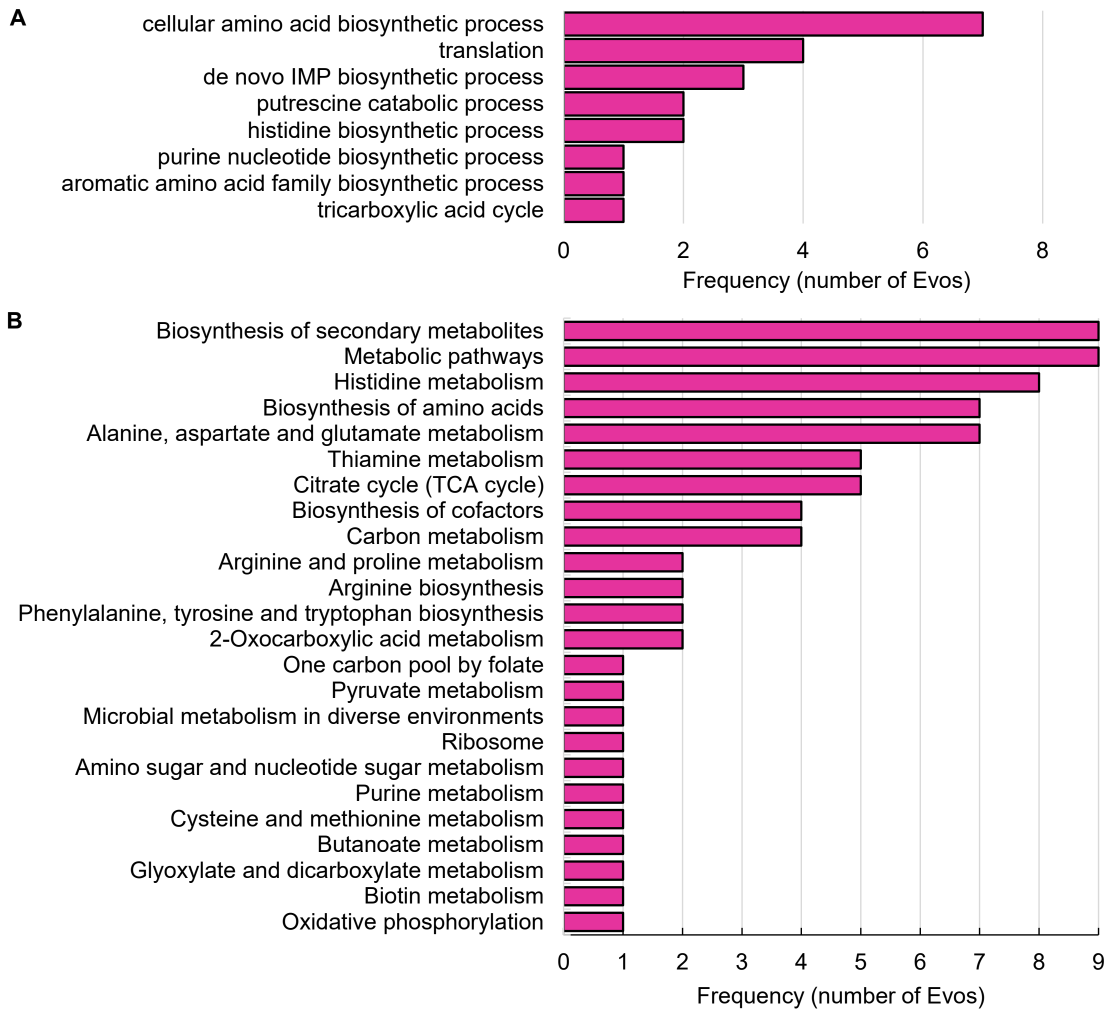


**Figure S5 Enriched functions of DEGs in Evos. A.** KEGG pathways. **B.** GO Terms. The numbers of the Evos sharing the same enriched functions are shown.

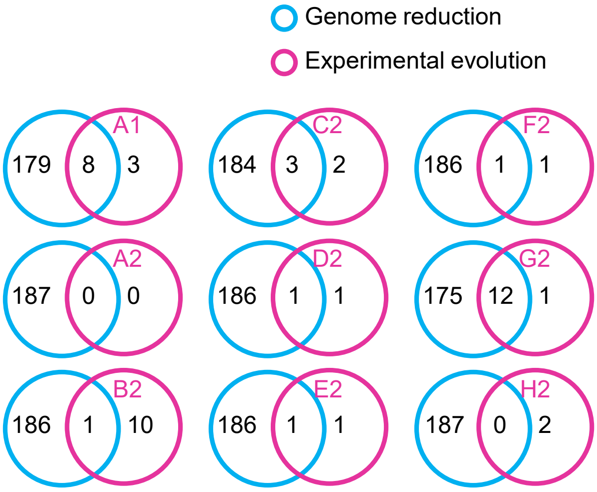


**Figure S6 Venn diagrams of the DEGs determined by RankProd**. The numbers of individual and overlapped DEGs induced by genome reduction and evolution are indicated.

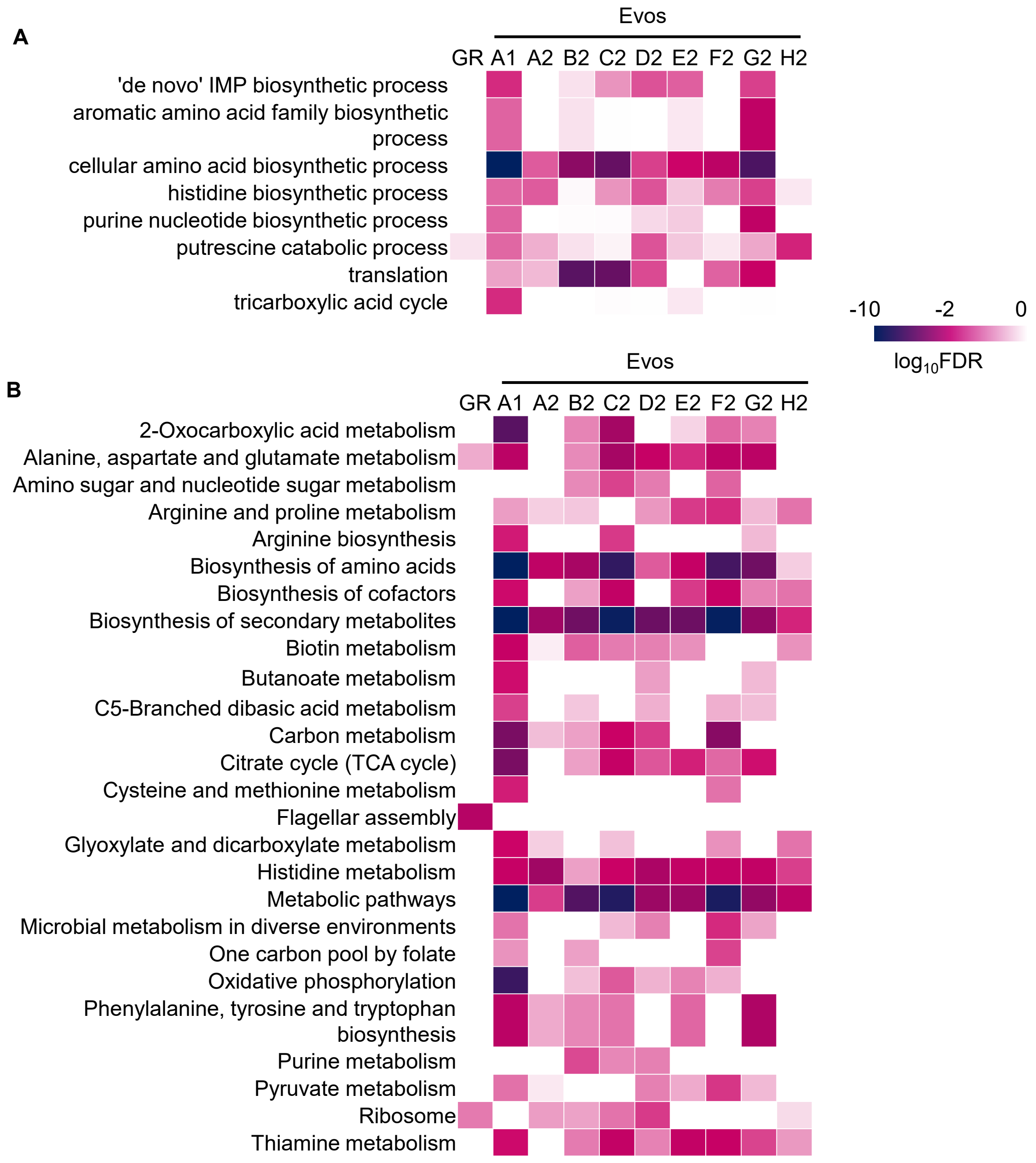


**Figure S7 Heatmap of the significance of enriched functions. A.** KEGG pathway. **B.** GO term. GR and Evos indicate the genome reduction and evolution, respectively. The statistical significance (FDR) of the enriched pathways and biological processes is shown on a logarithmic scale represented by color gradation.

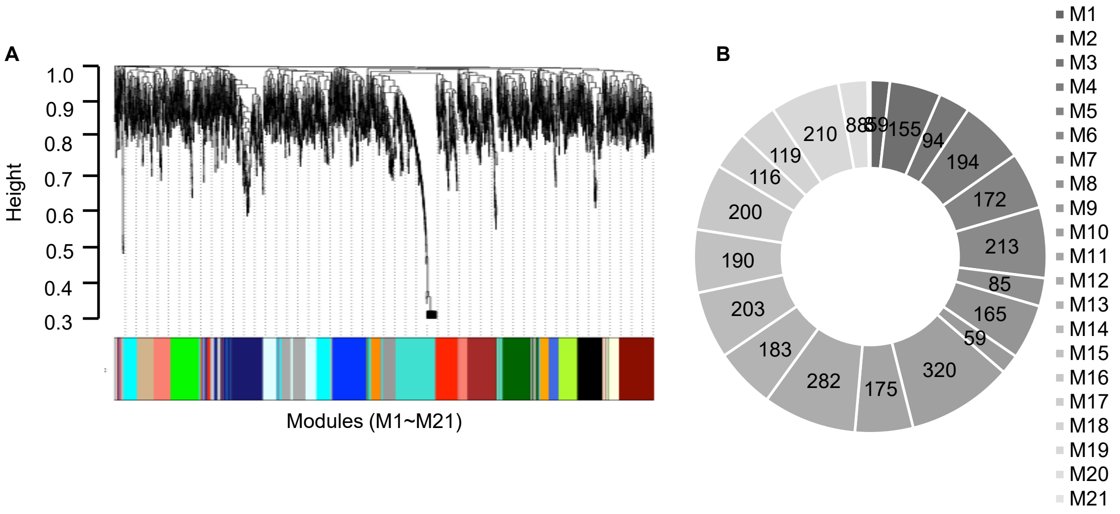


**Figure S8 Reconstruction of gene modules according to transcriptomes. A.** Gene modules clustered by weighted gene co-expression network analysis (WGCNA). **B.** Gene modules reconstructed by WGCNA. A total of 21 modules newly constructed are represented in gradation. The number of genes assigned in each module is indicated.

**Table S1 Daily records of experimental evolution.** The time and $OD_{600}$ of overnight culture, the well (dilution rate) used for the serial transfer, and the calculated generation and growth rate are shown. All nine evolutionary lineages are summarized.

**Table S2 Statistics of genome sequencing.** The parameters acquired in the genome resequencing, which represent the goodness of the sequencing, are summarized.

**Table S3 List of genome mutations.** The total 65 mutations fixed in the Evos are summarized. The type of mutation, the position in the reduced genome, distance to the nearest genomic scar, changes in DNA and amino acid, gene function, essentiality (e, essential; n, nonessential), gene category, etc., are indicated. Note that the entire population held the mutations, i.e., 100% frequency in DNA sequencing.

**Table S4 Statistics of chromosomal periodicity of transcriptomes.** The maximal peak (wavelength) acquired by the Fourier transform, the number of periods resulting from the curve fitting, the Fisher-g test and the p values are summarized.

**Table S5 List of the overlapped DEGs.** A total of 108 DEGs are summarized. Gene ID, gene name, and gene function are indicated.

**Table S6 Datasheet of normalized gene expression.** Gene expression levels are shown in the logarithmic value of FKPM. Gene ID, gene name, strain name, and nine evolutionary lineages are indicated. N0 and N28 represent the wild-type and reduced genomes, respectively.
